# Exposure to hypergravity during zebrafish development alters cartilage material properties and strain distribution

**DOI:** 10.1101/2020.05.26.116046

**Authors:** Elizabeth A Lawrence, Jessye A Aggleton, Jack J. W. A. van Loon, Josepha Godivier, Robert L. Harniman, Jiaxin Pei, N. C. Nowlan, Chrissy L Hammond

## Abstract

Terrestrial vertebrates have adapted to life on Earth and its constant gravitational field, which exerts load on the body and influences the structure and function of tissues. While the effects of microgravity on muscle and bone homeostasis are well described, the effects of shorter exposures to increased gravitational fields are less well characterized. Here, we exposed zebrafish to 3 and 6g hypergravity from 3-5 days post fertilisation, when key events in jaw cartilage morphogenesis occur. We did not observe changes to growth, or morphology of cartilage or muscle. However, we observed altered mechanical properties of jaw cartilages. We model the impact of these material property changes using Finite Element Analysis and show strain distribution in the jaw is altered following hypergravity. In regions of predicted altered strain we observed local changes to chondrocyte morphology, suggesting altered gravity affects chondrocyte maturation, ultimately leading to changes to cartilage structure and function.

## Background

Mechanical loading of the skeleton occurs during physical activity through muscle contraction and ground reaction forces (Lanyon *et al.*, 1975; Usui *et al.*, 2003). This loading builds and maintains bone mass, making increased skeletal loading an area of interest in the treatment of osteoporosis (Russo and MD, 2009). A physiological level of mechanical loading is beneficial to cartilage function *in vitro* by increasing chondrocyte proliferation and anabolic processes, boosting proteoglycan synthesis (Klein-Nulend *et al.*, 1987; Lee and Bader, 1997; Soltz *et al.*, 2000; Shelton, Bader and Lee, 2003; Sharma, Saxena and Mishra, 2007). *In vivo* experiments in hamsters (Otterness *et al.*, 1998), rats (Galois *et al.*, 2003), and humans (Manninen, 2001), have indicated that moderate exercise has a chondroprotective role, resulting in decreased risk of severe osteoarthritis (OA). In contrast, overloading or reduced loading of joints has a role in cartilage destruction by promoting catabolic pathways. Extreme loading (through high impact sports (Arendt and Dick, 1995; Levy *et al.*, 1996) or joint misalignment (Meireles *et al.*, 2017)) leads to extracellular matrix damage, loss of collagen, chondrocyte cell death and eventually OA (Torzilli *et al.*, 1999; Loening *et al.*, 2000; Patwari *et al.*, 2004).

As mechanical loading is exerted on the skeleton by gravitational forces (Kohrt, Barry and Schwartz, 2009), many studies on the musculoskeletal system have been carried out across a range of gravity levels. Microgravity has significant, well documented, effects on the skeleton with decreased bone density observed in humans (Demontis *et al.*, 2017) and fish (Chatani *et al.*, 2015), and disruptions to skeletal maturation observed in immature mice (Maupin *et al.*, 2019). Studies on astronauts and cosmonauts following long duration space flight found that 92% had a minimum of 5% bone loss in at least one skeletal site (LeBlanc *et al.*, 2007) with weight-bearing regions most affected (LeBlanc, Shackelford and Schneider, 1998; Vico *et al.*, 2000). Sarcopenia is also observed in microgravity, with decrease in muscle volume of around 15% following 4-6 months in microgravity (Adrian LeBlanc *et al.*, 2000). While decreased mechanical loading in microgravity has been uniformly associated with disuse bone loss, exposure to hypergravity has been shown to increase or decrease bone depending on the degree of hypergravity. One study exposed mice to hypergravity for 21 days and found that at 2g, there was an improvement in trabecular bone volume, fewer osteoclasts and an increase in mineralization (Gnyubkin, 2015). At 3g they found cortical thinning, more osteoclasts and a reduced rate of bone formation (Gnyubkin, 2015), supporting the idea that loading is beneficial to a point, after which it becomes deleterious (Yokota, Leong and Sun, 2011). Zebrafish larvae have been exposed to hypergravity in a Large Diameter Centrifuge (LDC). Aceto et. al. (2015) exposed zebrafish to 3g and observed increased ossification in the cranial skeleton of larvae exposed to 3g between 5 and 9 days post fertilisation (dpf) (Aceto *et al.*, 2015).

Another component of the musculoskeletal system sensitive to alterations in biomechanics is cartilage, particularly the articular cartilage of synovial joints. This cartilage has a limited regenerative capacity (Karuppal, 2017) and is important for absorbing load to protect the underlying bone, ensuring the smooth function of joints (Fox, Bedi and Rodeo, 2009). Human bed-rest studies, hind-limb unloading studies in rats and studies performed on mice exposed to real microgravity have demonstrated that loss of mechanical forces lead to cartilage degradation primarily through proteoglycan loss (Souza *et al.*, 2012; Ganse *et al.*, 2015; Willey *et al.*, 2016; Fitzgerald *et al.*, 2019). Cell culture experiments carried out in microgravity also support the observation of cartilage degradation under reduced loading conditions, with cytoskeletal reorganization and extracellular matrix (ECM) composition altered following short exposures (Van Loon *et al.*, 1995; Freed *et al.*, 1997; Zhang *et al.*, 2003; Ulbrich *et al.*, 2010; Aleshcheva *et al.*, 2013). In comparison to work on cartilage in unloading conditions, less is known about the effect of hypergravity and increased mechanical loading. One study on cultured chondrocytes showed downregulation of *BMP4* (crucial in collagen type II and aggrecan synthesis (Reddi, 2001)) following very short term cyclic hypergravity exposure during parabolic flight (Wehland *et al.*, 2015), suggesting that articular cartilage health would be impaired under such loading conditions.

Here, we show that exposure to hypergravity for 48 hours from 3 days post fertilisation (dpf) in zebrafish has no substantial effect on craniofacial cartilage morphology or musculature, but causes significant changes to cartilage material properties, chondrocyte morphology and ECM organisation. We also demonstrate altered strain distribution across the lower jaw following hypergravity exposure, providing an explanation for the cell-level changes. Altogether, this shows that hypergravity exposure in zebrafish larvae between 3-5dpf can induce subtle, but detrimental, changes to cartilage which could become more severe over time.

## Materials and methods

### Zebrafish husbandry and mutant lines

Zebrafish were maintained as described previously (Westerfield, 2000). Experiments were approved by the European Space Agency (ESA) and performed in accordance with UK ASPA regulations.

### Hypergravity experiments

Zebrafish were exposed to hypergravity in the Large Diameter Centrifuge (LDC) (Van Loon *et al.*, 2008) at the European Space Research and Technology Centre (ESTEC) for 48 hours from 3dpf to 5dpf. The LDC consists of a central axis linked to 2 arms. Samples can be placed in 6 gondolas (which can be set to 2 different hypergravity levels) plus 1 central gondola at 1g to control for rotation and possible related Coriolis accelerations (Van Loon, 2007). The larvae were exposed to 3g and 6g, with control larvae located at the central axis, further larvae were maintained at 1g static (Supplementary Figure 1A). In each gondola, 4 petri dishes containing 150ml of Danieau’s solution and <35 larvae each were stacked in the centre of an incubator set to a constant temperature of 28°C (Supplementary Figure 1A-D). Larvae were incubated in the dark except during the recording of videos (to monitor survival and swim behaviour during the experiment). Following exposure to hypergravity, 1 petri dish from each gravity condition was reserved for behavioural studies, with the rest fixed in 4% paraformaldehyde (PFA) or bone fix (3.5% formaldehyde in 40mM phosphate buffer) for further analysis.

### Whole fish measurements

Larvae were mounted in glycerol and imaged on a Leica MZ10F stereo microscope. Head to tail length was measured using the line function in Fiji (Schindelin *et al.*, 2012) (Supplementary Figure 1E).

### Antibody labelling

Immunohistochemistry was performed as previously described (Hammond and Schulte-Merker, 2009). Briefly, larvae were fixed in 4% PFA and dehydrated to 100% MeOH for storage, then rehydrated to 1 x PBST, permeabilised and blocked in 5% horse serum prior to incubation in primary antibodies (Collagen type II [Abcam ab34712] 1:500, A4.1025 [Developmental Studies Hybridoma Bank] 1:500, L-plastin (Cvejic *et al.*, 2008) 1:200, Acetylated tubulin [Sigma-Aldrich T6793] 1:200) at 4°C overnight (o/n). Samples were washed three times in PBST, re-blocked and incubated in secondary antibodies (Goat anti-rabbit 555 [Dylight], Donkey anti-mouse [Thermofisher Scientific], Goat anti-chick 647 [Thermofisher scientific] all at 1:500) o/n at 4°C. Where required, larvae were incubated in 5μg/ml 4′,6-diamidino-2-phenylindole (DAPI) [Invitrogen] for 1 hour and washed in PBST prior to imaging. Samples were mounted in 0.5% low melting point agarose and imaged on a Leica SP5II confocal microscope with a 10x or 20x objective.

### Alcian blue and Alizarin red staining

Alcian Blue and Alizarin red whole mount larval staining was performed as previously described (Walker and Kimmel, 2007) on larvae fixed in 3.5% formaldehyde.

### Atomic force microscopy

Atomic Force Microscopy (AFM) was conducted utilizing a Multi-mode VIII microscope with Nanoscope V controller, operating in a PeakForce control regime [Bruker, CA, USA]. Larval cartilage was investigated in a hydrated state in an ambient environment (Elizabeth A. Lawrence *et al.*, 2018). Prior to AFM investigation, larvae were fixed in 4% formaldehyde, stained with Alcian blue and Alizarin red (as above), and the lower jaw was dissected in 1% glycerol in PBS to prevent structural changes induced by drying. For measurement of Young’s modulus (YM) via Quantitative Nano-mechanical Mapping (QNM) RTESPA-150 cantilevers [Bruker, Ca, USA] were utilized, having nominal spring constant and tip radii of 5 N/m and 8 nm respectively. Cantilevers were calibrated via the relative method utilizing a PDMS standard with data fit to a DMT model accounting for the effect of adhesion forces in the standard Hertzian model for indentation. Three fish were investigated for each level of hypergravity exposure. For each fish, three separate 500nm x 500nm regions were scanned in both the immature chondrocytes and hypertrophic chondrocytes, six regions in total. 65,536 measurements were taken per scanned region and their RMS average recorded for comparison. Data was normalised to values from 1g static samples to show the relative YMs (Melbourne *et al.*, 2018; Nigmatullin *et al.*, 2018, 2019; Swift *et al.*, 2018; Terry *et al.*, 2019; Gubała *et al.*, 2020).

### Nanoindentation

Larvae were fixed in 4% PFA and stored in 100% MeOH before rehydration to 30% sucrose in PBS. Samples were submerged in 30% sucrose in PBS, diluted 1:2 in optimum cutting temperature compound (OCT) until they sunk. This solution was refreshed for embedding and samples were flash frozen and sectioned in a coronal orientation using an NX70 Cryostat [ThermoFisher] at a thickness of 10µm. Nanoindentation was performed on sections containing the jaw joint and/or Meckel’s cartilage using a Chiaro nanoindentation device [Optics11, Amsterdam, The Netherlands]. Sections were kept submerged in PBS at room temperature whilst measurements were taken. A spherical nanoindentation probe with an 8µm radius and stiffness of 0.49N/m was used, and tissues were indented to a depth of 1µm with velocity of 1µm s^−1^, with the tip held at a constant depth for 10s. The collected curves were analysed based on Hertzian contact theory for direct comparison with AFM measurements and the resultant YM E_hertz_ were calculated assuming sample’s incompressibility. Nanoindentation was performed across all sections containing the joint or Meckel’s cartilage, with one measurement collected per region of interest in each section. The resulting YM were averaged for each region across sections. Nanoindentation was performed on five fish from each of the 1g static and 6g spin groups.

### Histological staining

Fixed samples were processed into paraffin, cut in 5µm sections, deparaffinised and stained with Haematoxylin & Eosin (H&E) and Alcian blue, Picrosirius Red, Safranin O/Fast Green or Masson’s Trichrome. ***H&E Alcian blue*** slides were stained in Erhlic’s haematoxylin for 5 minutes, rinsed, differentiated in acid alcohol and Scotts water, placed in eosin solution for 10 seconds, rinsed and immersed in Alcian blue for 30 minutes. ***Picrosirius Red*** slides were immersed in picrosirius red for 1 hour and washed in 2 changes of acidified water. ***Safranin O***/***Fast Green*** slides were stained with Weigert’s iron haematoxylin for 10 minutes, washed for a further 10 minutes, stained with 0.05% fast green solution for 5 minutes, rinsed in 1% acetic acid and immersed in 0.1% Safranin O solution for 5 minutes. ***Masson’s Trichrome*** Sections were re-fixed in Bouins solution, stained in Weigert’s iron haematoxylin then Ponceau Fuschin (Masson’s) for 5 minutes, rinsed, immersed in phosphomolybdic acid solution and counter stained with Aniline blue both 5 minutes and dipped in 1% acetic acid for 10 seconds. Following staining, all sections were dehydrated sequentially to 100% industrial methylate spirit (IMS), immersed in xylene 3 x 5 minutes and mounted using DPX mountant. Slides stained with Picrosirius red were imaged under polarized light on a Leica DMI6000 inverted epifluorescence microscope, all other slides were imaged on Leica MZ10F stereo microscope.

### Measurement of staining intensity from histology slides

To measure the staining intensity of Safranin O and Masson’s trichrome, images were opened in Fiji and the segmented line tool was used to draw a line through the ECM surrounding immature and hypertrophic chondrocytes in the jaw region. The plot profile command was then performed to extract the gray value along this line and measurements normalised to the image background to remove white balance discrepancies. This was performed in ten areas of immature and hypertrophic chondrocytes respectively, with measurements recorded from three fish per gravity condition.

### Cell circularity and area quantification

Chondrocyte morphology was measured in Fiji from brightfield images of alcian blue stained lower jaw paraffin sections (5µm thick). The polygon selection tool was used to outline chondrocytes at the joint and intercalation zone of the Meckel’s cartilage (Figure 2 A), the area and roundness shape descriptors were collected using the measure command. Cells from the middle of the Meckel’s cartilage were classified as hypertrophic and cells from the jaw joint and Meckel’s symphysis (black asterisk in Figure 2 A) were classified as immature.

### Jaw measurements

Confocal image stacks of the lower jaw immunostained for type II collagen were loaded in to Fiji (Schindelin *et al.*, 2012) and the line tool, followed by the measure command, were used to take length and width measurements (Figure 1 A). 3D jaw renders and joint measurements were executed as previously described in (Elizabeth A. Lawrence *et al.*, 2018) (Figure 1 D) using Amira 6.0 [FEI].

**Figure 1:**
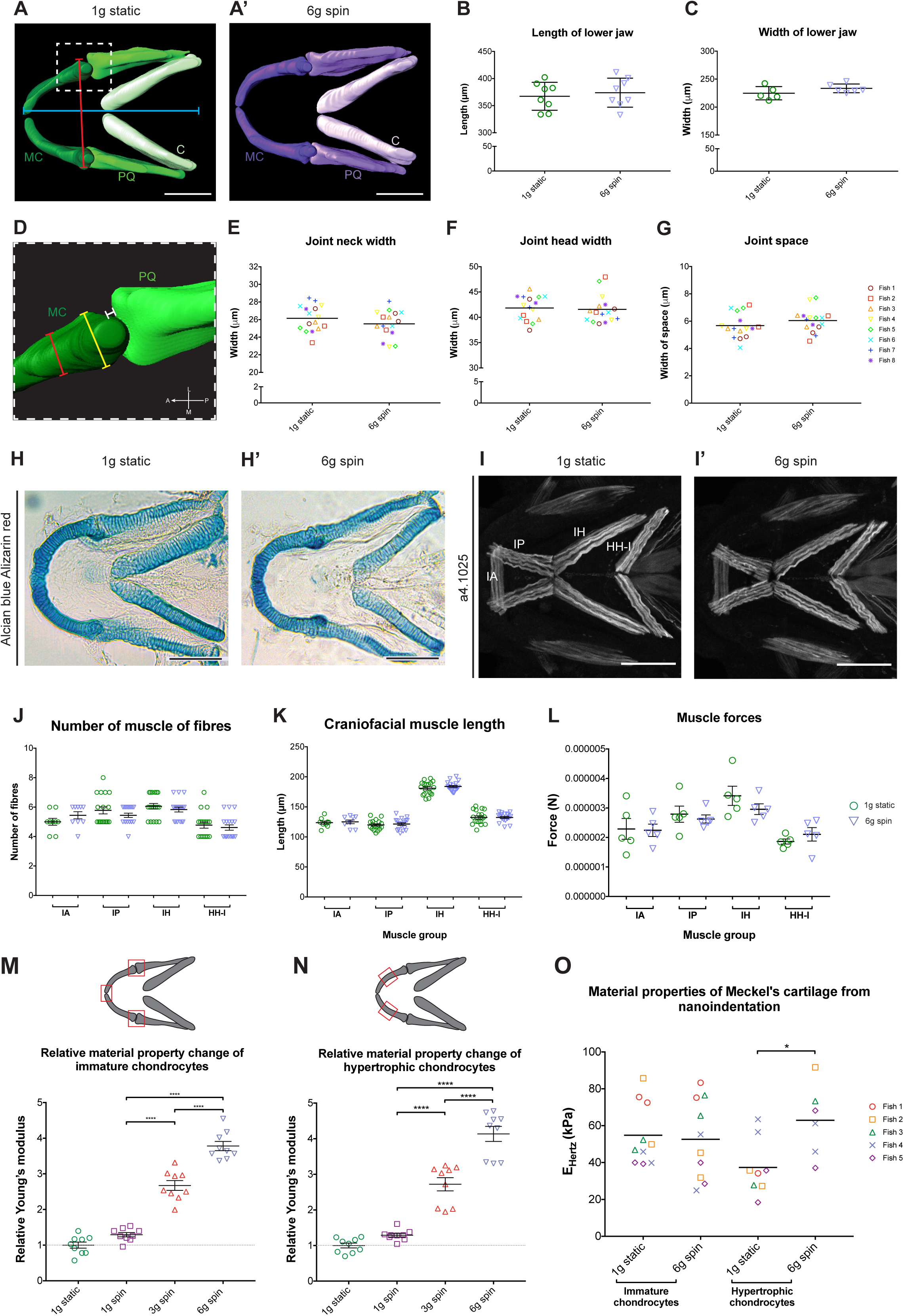
Lower jaw morphology and musculature are unchanged following hypergravity exposure, but changes to cartilage material properties are observed. (A, A’) 3D surface renders from confocal image stacks of lower jaw cartilage in ventral orientation from 1g static (A) and 6g (A’) zebrafish at 5dpf (MC = Meckel’s cartilage, PQ = palatoquadrate, C = ceratohyal). Scale bar: 100μm. (B, C) Quantification of lower jaw length (B) and width (C). Location of measurements shown by red (width) and blue (length) line in (A) (n=8 for all, different symbols = individual fish). (D) Close-up image of jaw joint from 1g static 3D render, position in lower jaw shown by white dashed box in (A). Orientation compass: A, anterior; L, lateral; M, medial; P, posterior. (E-G) Quantification of joint neck (E) and joint head (F) width, and joint space (G), location of measurements shown in (D), red line = joint neck, yellow line = joint head, white line = joint space (n=8 for all). (H, H’) Brightfield images of Alcian blue Alizarin red stained lower jaws from 1g static (H) and 6g (H’) conditions. Scale bar = 100μm. (I, I’) Maximum projections of ventral confocal image stacks from 5dpf 1g static (I) and 6g (I’) zebrafish immunostained for myosin (A4.1025) (IA = Intermandibularis anterior, IP = Intermandibularis posterior, IH = Interhyoideus, HH-I = Hyohyoideus inferior). Scale bar = 100μm. (J, K) Quantification muscle fibre number (J) and length (K) measured from confocal image stacks. Location of muscle groups shown in (I). (L) Quantification of craniofacial muscle forces. (M, N) Relative YM values from AFM for immature (M) and hypertrophic (N) chondrocytes from 1g static, 1g, 3g and 6g zebrafish (n=3 for all). Location of measurements taken shown in schematic above graphs. (O) Material properties determined by nanoindentation in 1g static and 6g zebrafish. Data is mean with s.e.m. (E-G show mean with no s.e.m.), D’Agostino and Pearson test performed for all data, followed by student’s unpaired t-test in B, C, E, F and G. One-way ANOVA performed within muscle groups in J, K and L and in M and N. Mann-Whitney u-test in O. * p ≤ 0.05, ** p ≤ 0.01, *** p ≤ 0.001, **** p ≤ 0.0001.

### Muscle quantifications and calculation of muscle forces

Muscle forces for each muscle group were calculated using methodology from (Brunt *et al.*, 2016). In brief, muscle fibre number and length was quantified manually in Fiji from confocal images of A4.1025 stained zebrafish and the cross-sectional area of the fibers was calculated using the formula: *πr*^*2*^. The radii of the fibres was calculated by taking a measurement across each fibre and dividing it by two. To calculate the cross-sectional area of the whole muscle group, the resulting value was multiplied by the number of muscle fibers. This area was multiplied by the maximal force generated per unit area for larval zebrafish skeletal muscles (40n N/µm^2^, from (Iorga *et al.*, 2011)) to give the final force value for the muscle.

### Finite Element Analysis

Finite Element (FE) models of the lower jaw for 1g static and 6g conditions were created from a confocal image stack of a representative experiment specimen. The same cartilage morphology and muscle forces were used for both conditions’ models in the absence of significant differences in cartilage and muscle morphology between 1g static and 6g specimens (Figure 1A-C, H, H’). The FE meshes were developed and modelled for jaw opening and closing movements as previously published (E.A. Lawrence *et al.*, 2018). Two different versions of the model were created, one for the nanoscale properties derived from AFM and the other using the microscale properties measured by nanoindentation. Relative material properties were derived from AFM and nanoindentation experiments (values used are listed in Table 1). For AFM and nanoindentation experiments separately, values were normalised relative to the 1g static hypertrophic chondrocyte Young’s modulus. In both experiments the Young’s modulus of the joint interzones were set at 0.025% of the 1g static hypertrophic chondrocyte Young’s modulus, a markedly lower value than the chondrocyte cells to enable realistic joint movement, and a Poisson’s ratio of 0.495. To reflect the respective experimental Poisson’s ratios, in the nanoindentation experiment hypertrophic and immature chondrocytes Poisson’s ratios were set at 0.495; in the AFM experiment hypertrophic and immature chondrocytes Poisson’s ratios were set at 0.3. All muscle forces were calculated based on cross-sectional area of the anatomical muscles, with forces reduced to 60% of maximal force (to represent jaw respiratory movement).

**Table 1:** Actual and relative material property values of immature and hypertrophic chondrocytes from AFM and nanoindentation which were used for FE models. Values represent the mean measurement across samples and the figure number of the corresponding FE model is shown in the right-hand column.

| Gravity condition | Method | Hypertrophic chondrocytes |  | Immature chondrocytes |  | Figure no. of corresponding model |
| --- | --- | --- | --- | --- | --- | --- |
|  |  | Actual material property value | Relative material property value | Actual material property value | Relative material property value |  |
| 1g static | AFM | 7.7 MPa | 1 | 4.2 MPa | 0.51968 | Figure 3 A |
|  | Nanoindentation | 37.39 kPa | 1 | 54.8 kPa | 1.46563 | Figure 3 B |
| 6g spin | AFM | 31.0 MPa | 4.13444 | 15.2 MPa | 1.99503 | Figure 3 A' |
|  | Nanoindentation | 62.9 kPa | 1.68227 | 52.63 kPa | 1.407597 | Figure 3 B' |

**Table 2:**
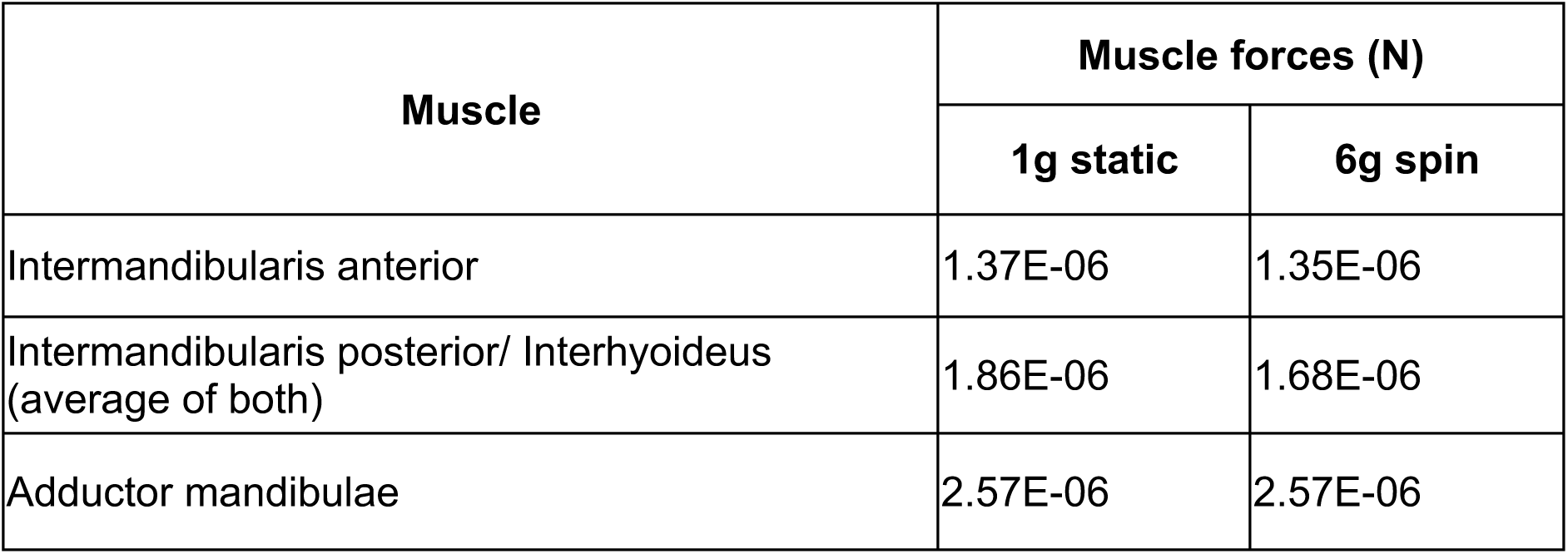
Muscle forces used for FE model generation. Values for 1g static and 6g spin fish represent 60% of the maximum muscle force calculated for each muscle group in Figure 1 L with the exception of the adductor mandibulae.

### Transmission Electron Microscopy

Transmission electron microscopy (TEM) was performed on 5dpf larvae fixed in 2.5% glutaraldehyde in 0.1M sodium cacodylate for 1 hour at RT. These samples were embedded in 3% agarose before being osmium/uranyl acetate stained, dehydrated and infiltrated with Epon in a Leica EM TP tissue processor using the standard protocol. Prior to sectioning, samples were laterally embedded in 100% Epon and left to harden at 60°C for two days. These blocks were sectioned at a thickness of 70nm on a Leica EM UC7 RT ultramicrotome using a diamond knife [Diatome]. Sections were dried o/n before staining in uranyl acetate for five minutes followed by dH2O washes, five minutes in lead citrate and a final dH2O wash. Sections were imaged on a Tecnai 12 -FEI 120kV BioTwin Spirit Transmission Electron Microscope.

### Analysis of collagen fibre density from TEM images

To analyse collagen fibre density, TEM images were loaded into Fiji and a region of interest (ROI) of 1m^2^ was drawn in a random location containing ECM. The number of collagen fibres in this ROI was counted using the multipoint tool and this process was repeated for five separate regions per image. Four images were analysed from two separate sections per fish, with the sections originating from one fish per gravity condition.

## Results

### Craniofacial cartilage morphology and musculature are unaffected by hypergravity

Having confirmed there was no delay in larval growth (Singleman and Holtzman, 2014) following hypergravity exposure (Supplementary Figure 1E), we investigated the effect of increased mechanical loading through hypergravity on developing cartilage, using type II collagen immunostaining, and Alcian blue and Alizarin red double staining to visualize morphology (Supplementary Figure 2A, Figure 1A, A’, H, H’). These analyses did not reveal significant changes to the overall jaw shape, with jaw length and width not significantly changed in 1g static and 6g fish (Figure 1B, C). Analysis of joint morphology also revealed no significant difference in fish exposed to hypergravity compared to the 1g controls (Figure 1 - G).

Given the association of microgravity with muscle loss (Martin, Edgerton and Grindeland, 1988; Caiozzo *et al.*, 1994), we stained larvae with the pan-skeletal myosin marker A4.1025 to visualize muscles in the lower jaw (Figure 1I, I’,). From these images we quantified muscle fibre number, length and force, with no significant differences in craniofacial muscle seen between zebrafish incubated in normal gravity and at 6g between 3-5 dpf (Figure 1J, K, L).

### Material properties are altered in the lower jaw

Changes to mechanical loading have been observed to change skeletal stiffness at both at the nano and micro scales (Turko *et al.*, 2017). This led us to investigate the relative material properties of lower jaw cartilage using AFM (Melbourne *et al.*, 2018; Nigmatullin *et al.*, 2018, 2019; Swift *et al.*, 2018; Terry *et al.*, 2019; Gubała *et al.*, 2020). Measurements were taken from areas of immature chondrocytes and hypertrophic chondrocytes (location of measurements shown by schematics in Figure 1M, N). In both instances, a positive correlation between the magnitude of gravitational exposure and the measured YM was seen. Fish from 3g and 6g had a significantly higher YM than 1g static or 1g spin fish (Figure 1M,N)), with 6g showing a significant increase in YM compared to 3g fish. This trend represents a stiffening of the cartilage following hypergravity exposure during development.

Within complex materials, different structures can have a greater influence on stiffness at different length scales. Having used AFM to measure YM at the nano-scale we employed nanoindentation to investigate material properties of lower jaw cartilage at the micro-scale. Measurements from nanoindentation show 6g fish had a significantly higher YM (E_hertz_) in hypertrophic chondrocytes when compared to 1g static fish. This pattern of increased stiffness was not seen for immature chondrocytes (Figure 1O).

### Exposure to hypergravity affects chondrocyte maturation and behaviour

The impact of hypergravity at a cellular level was evaluated by measuring chondrocyte morphology from Alcian blue stained lower jaw sections (Figure 2A - C). Chondrocyte area was significantly reduced in immature and hypertrophic regions in 6g fish, with immature cells at the joint and Meckel’s symphysis showing the largest area reduction (Figure 2B). Immature chondrocytes also showed a significant decrease in circularity (Figure 2C). The regions of most change to cell morphology were co-localised to muscle attachment sites in the mid-Meckel’s cartilage and at the jaw joint. This suggests that short exposure to hypergravity alters chondrocyte behaviour, causing cartilage and resulting bones to develop abnormally if maintained in hypergravity for an extended period.

**Figure 2:**
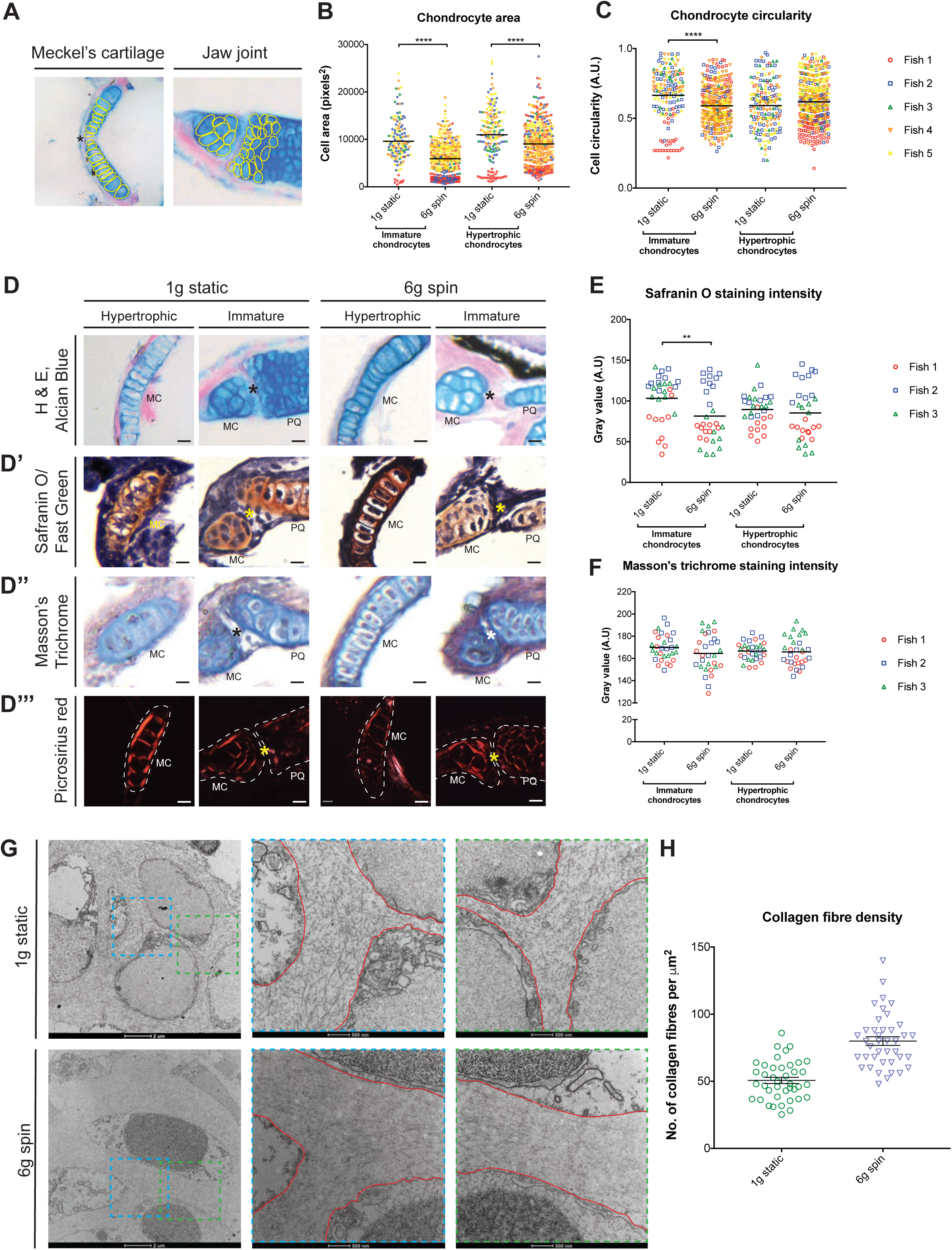
Subtle changes to chondrocytes, and their surrounding ECM occur in areas of altered strain. (A) Representative image with chondrocytes included in area and circularity measurements outlined in yellow. Black asterisk = Meckel’s symphysis. (B, C) Quantification of chondrocyte area (B) and circularity (C) (n=5 for all, colours = individual fish). (D-D’’’) Alcian blue, haematoxylin and eosin (D); Safranin O and fast green, (D’); Masson’s trichrome (D’’); and picrosirius red (D’’’) stained ventral sections in 1g static and 6g fish. Asterisk = centre of joint, dotted line in (D’’’) = outline of cartilage element from section. Scale bar = 10μm. MC = Meckel’s cartilage, PQ = palatoquadrate, C = ceratohyal. (E, F) Quantification of Safranin O (E) and Masson’s trichrome (F) staining intensity. (G) Electron micrographs of hypertrophic chondrocytes in 1g static and 6g zebrafish. Dashed areas = higher magnification images displayed in the centre and to the right of the panel, red lines = chondrocyte borders. Scale bars are shown below each image. (H) Quantification of collagen fibre density in the ECM of 1g static and 6g fish (n = 1 for both). All data is mean with s.e.m. D’Agostino and Pearson normality test performed in B, C, E and F: followed by Mann-Whitney u-test in B and C, and student’s unpaired t-test in E and F. * p ≤ 0.05, ** p ≤ 0.01, *** p ≤ 0.001, **** p ≤ 0.0001.

### Histological staining reveals subtle changes to the ECM surrounding chondrocytes in areas of altered cell morphology

As increased mechanical loading is associated with higher glycosaminoglycan (GAG) synthesis and decreased extracellular matrix (ECM) secretion (Schröder *et al.*, 2019), we performed wholemount Alcian blue and Alizarin red double staining (Figure 1H, H’) and Alcian blue, H & E on sections to visualize GAGs throughout the lower jaw (Figure 2D). From brightfield images, we observed no changes to mineralisation or GAGs at 5dpf (Figure 1H, H’, 2D), suggesting secretion of this ECM component is unaffected by altered loading.

The effect of hypergravity exposure on other ECM components was examined using Safranin O, Masson’s trichrome, and picrosirius red staining on sections of the lower jaw including the joint and Meckel’s cartilage. Proteoglycan content and mineralisation of the cartilage ECM was visualized with Safranin O/ Fast Green staining which marks the cartilage red, according to the amount of proteoglycan present, and bone in green (Figure 2D’). The intensity of this stain was measured, revealing fish from the 6g condition had a significantly lower staining intensity in regions of immature and hypertrophic chondrocytes than 1g static fish (Figure 2E), corresponding to a reduction in proteoglycan content in the cartilage following hypergravity exposure. This reduction was more pronounced in regions of ECM surrounding immature chondrocytes (Figure 2E). No areas of mineralisation were seen in the stained sections so no information on bone formation could be gathered from this technique.

Masson’s trichrome staining was used to test whether hypergravity impacted collagen content in chondrocyte ECM (Figure 2D’’). This stain shows collagen in blue, and measurements of staining intensity showed no significant change to collagen content in the ECM surrounding immature or hypertrophic chondrocytes in 6g fish compared to the ECM in 1g static fish (Figure 2F). Alongside measuring total collagen in the ECM through Masson’s trichrome, picrosirius red staining was used to assess the balance of type I and type III collagen fibers in the ECM. Under polarized light, ECM surrounding immature and hypertrophic chondrocytes appeared red/orange (Figure 2D’’’), indicating a predominance of type I collagen fibres over type III (which would give green birefringence). This was unchanged in zebrafish from the 6g condition (Figure 2D’’’).

### Hypergravity causes changes to collagen fibre packing in the ECM

To further explore how the cartilage ECM is affected by hypergravity, TEM was carried out on sections of ear cartilage, which are comparable to regions of hypertrophic chondrocytes in the lower jaw where the most significant change to material properties was seen. From the micrographs, subtle changes to the collagen fibre packing were observed (Figure 2G), with fibres appearing closer together in 6g fish. This observation was strengthened by quantification of fibre density, which revealed a trend for increased fibre density in 6g fish (Figure 2H).

Taken together with histology data, this suggests that hypergravity induces macromolecular changes in the cartilage which give rise to slight changes at the tissue level. If the larvae had been maintained in hypergravity for longer, it is likely that these changes would have become more pronounced, leading to more severe changes to the tissue.

### Finite Element Analyses reveal altered strain distribution in response to hypergravity exposure

To assess whether cell and matrix changes could be correlated with altered strain distribution in the lower jaw, FE models were generated. As no changes in lower jaw morphology or musculature were observed (Figure 1A-C, H, H’), the same volumetric model and muscle forces were used for both gravity conditions. Different normalised material properties from AFM (Figure 3A, A’) and nanoindentation (Figure 3B, B’) were applied to these models (Supplementary Table 1). From these models, it can be seen that maximum principal strain is more localised to the joint regions in 6g fish with stiffer cartilage, whereas in 1g fish strain is distributed over a larger area of cartilage elements (Figure 3A-B’). Both methods for obtaining material properties of the cartilage show similar change in strain pattern distribution following hypergravity exposure (from 1g static to 6g). Similarly, the pattern of von Mises stress is more evenly distributed throughout the jaw in the 1g static condition, whereas in the 6g condition stress is localised in regions already experiencing high stress, specifically the regions surrounding the joints (Supplementary Figure 3). This pattern is true for both opening and closing movements. The altered patterns of strain observed, in which the largest differences are close to the joint could provide an explanation for the subtle changes to cell maturation, which were more pronounced at the joint, and to changes observed to matrix packing.

**Figure 3:**
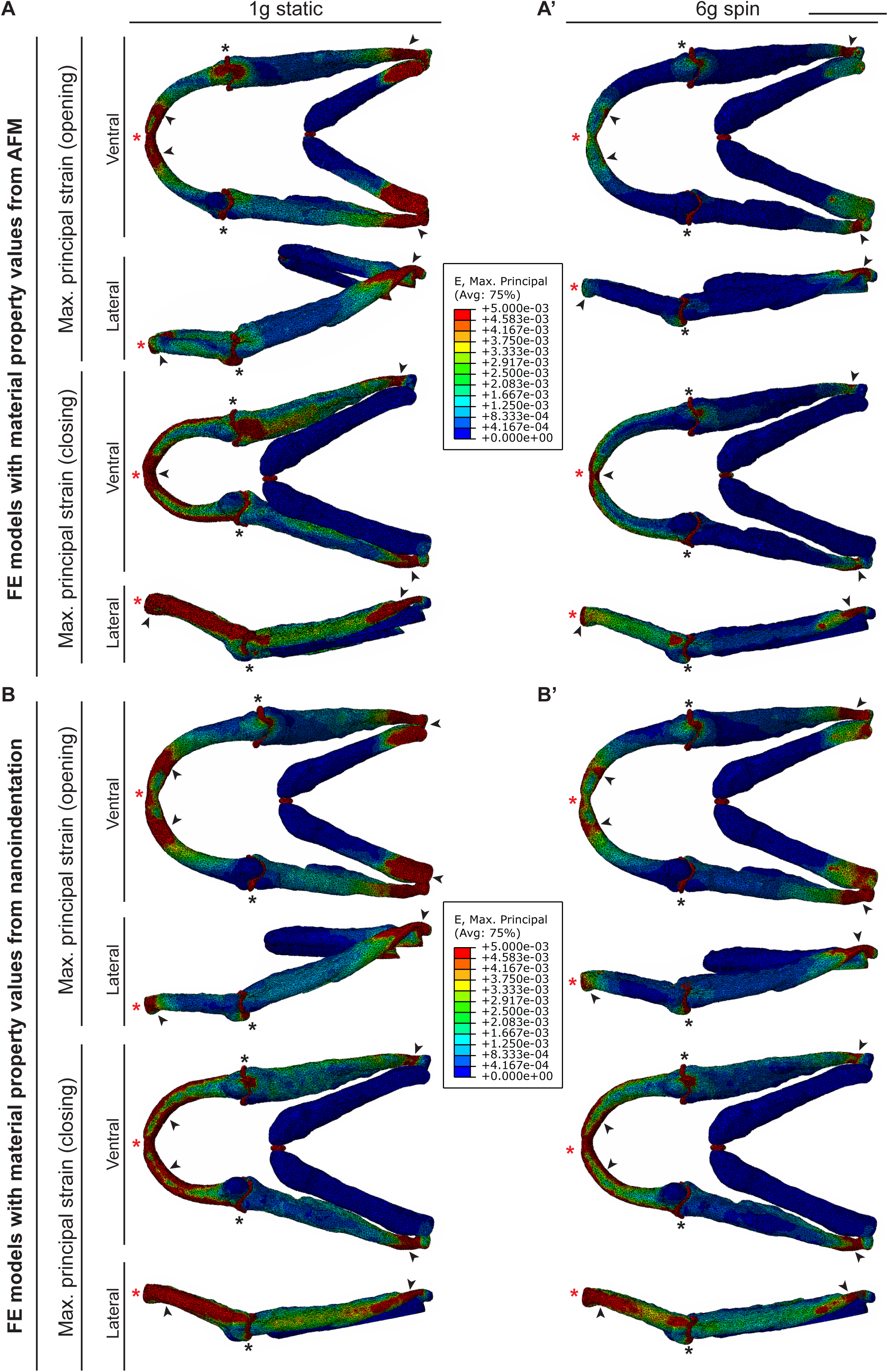
Altered ECM characteristics could result from altered strain distribution in the lower jaw following hypergravity exposure. (A,A’) FE models of maximum principal strain incorporating relative material property values from AFM in 1g static (A) and 6g zebrafish (A’). (B,B’) FE models of maximum principal strain incorporating relative material property values from nanoindentation in 1g static (B) and 6g zebrafish (B’). Black arrowheads = areas of high strain; black asterisks = jaw joints; red asterisks = Meckel’s symphysis. Ventral and lateral views shown for opening step in both gravity conditions.

## Discussion

The impact of microgravity on the musculoskeletal system has been well studied, with exposure to below-Earth gravity linked to muscle loss and decreased bone density. In comparison, relatively little is known about how short-term hypergravity exposure affects the musculoskeletal system. Here, we show that in zebrafish embryos gross cartilage and muscle morphology is unchanged, but cartilage material properties and its resulting biomechanical performance is affected by short-term exposure to hypergravity during development. We also show that hypergravity leads to altered chondrocyte maturation and subtle changes to the surrounding ECM, which may lead to more dramatic changes to the cartilage over time.

Gravity is important for cartilage health as it provides a loading force essential for cartilage homeostasis and prevention of degenerative diseases such as OA (Penninx *et al.*, 2001; Bader, Salter and Chowdhury, 2011; Musumeci, Szychlinska and Mobasheri, 2015; Mellor *et al.*, 2017). Although cartilage morphology and skeletal muscle mass has been shown to be affected by altered loading conditions (Vanwanseele *et al.*, 2003; Hinterwimmer *et al.*, 2004; Liphardt *et al.*, 2009; Gao *et al.*, 2018), our data suggest that two days of hypergravity exposure is not sufficient to cause gross morphological changes in larval zebrafish. One explanation for this is the length of exposure being insufficient to induce musculoskeletal remodeling, as previous studies have found that longer exposure to non-Earth gravity leads to more severe musculoskeletal transformations (A LeBlanc *et al.*, 2000; Demontis *et al.*, 2017). Another explanation for the lack of morphological alterations is that the level of hypergravity was not high enough to induce changes. However, it has previously been shown by Aceto et. al (Aceto *et al.*, 2015) that exposure to 3g between 5 and 9 dpf is sufficient to induce skeletal changes in zebrafish. This indicates that the age at which the zebrafish are exposed to hypergravity is also crucial (Franz-Odendaal and Edsall, 2018). Thus, we hypothesise that extending the amount of time larvae spend in hypergravity to include later time points key in musculoskeletal development would lead to more dramatic changes.

The hypothesis that more severe cartilage abnormalities would be seen following a lengthier hypergravity exposure is given weight by increased modulus seen in the matrix of zebrafish exposed to both 3g and 6g, with the largest increase seen in zebrafish subjected to 6g. FE analysis revealed that these altered material properties were sufficient to disturb strain pattern in the lower jaw, with the jaw joint and muscle attachment sites showing most change in the craniofacial cartilage elements. In these areas of altered strain, we observed abnormal chondrocyte maturation, which over time would likely give rise to altered joint shape and cartilage morphology. In conclusion short-term exposure to hypergravity in early development causes changes to ECM content and organisation in zebrafish which could induce more dramatic structural and morphological changes to the musculoskeletal system over extended periods of exposure.

## Acknowledgements

The authors would like to thank Kate Robson-Brown for her invaluable expertise, guidance and backing throughout the project. They would also like to express their gratitude to the ESA Education team, particularly Nigel Savage and Evelien Lageweg, and Alan Dowson at the LDC for their support and advice throughout the ‘Spin Your Thesis!’ campaign. Finally, they would like to thank David Labonte and Andrea Attipoe for the use of the Chiaro nanoindentation device and for their helpful assistance.

